# OmniT2I: a unified table-to-image framework for survival prediction and cross-modal inference with incomplete multimodal cancer data

**DOI:** 10.64898/2026.11.08.765158

**Authors:** Peng Li, Zuhair Majeed, Muhammad Khalid Khan Niazi, Merve Hasanov, Elshad Hasanov

## Abstract

Molecular profiles and histopathology provide complementary information about cancer prognosis, but these data are often incomplete for individual patients. Here we present OmniT2I, a table-to-image framework for survival prediction and cross-modal inference from incomplete multimodal cancer data. Each molecular profile was converted into a three-channel image representation with correlation-based feature placement. OmniT2I combines available molecular and histopathology embeddings through availability-aware fusion and predicts missing modalities from those observed. Across 10,373 patients with 32 cancer types, the C-index was 0.631 within cancer types and 0.783 with all cancers pooled (0.660 and 0.787 in 5,605 patients with all five modalities). A single model handled all 31 modality combinations and, compared with nine survival methods trained separately for each, achieved higher C-indices in 240 of 279 pooled and 226 of 279 within-cancer comparisons. Adding predictions of withheld modalities to observed inputs improved pooled C-indices in all 180 pairings of observed and withheld modality sets and within-cancer C-indices in 164. Adding measured histopathology to RNA produced the largest within-cancer C-index gain among the four additional modalities evaluated, whereas adding copy number variation reduced it. In seven external cohorts, frozen RNA-only C-indices differed from test performance in the corresponding cancer type in TCGA by a median of −0.003 (range, −0.092 to +0.079). OmniT2I enables prognostic modeling with incomplete multimodal data.

## Introduction

Cancer prognosis reflects molecular alterations, cellular programs and tissue organization. RNA, copy number variation (CNV), DNA methylation and protein abundance characterize different levels of molecular regulation, while hematoxylin and eosin (H&E) histopathology captures tumor morphology and its microenvironment. Their integration can connect these complementary observations to patient outcomes [1–3]. In practice, patients rarely have measurements from every modality. Restricting analysis to complete profiles discards information from patients with partial measurements, whereas fitting a separate model for each available combination fragments learning across overlapping patient groups. The scientific challenge is to learn prognostic relationships across molecular and tissue scales when the observations available for individual patients are incomplete.

Multimodal survival methods have addressed integration through shared representations [2–4], co-attention between histology and molecular features [5] and interactions between histology and biological pathways [6]. MultiSurv accommodates absent modalities [7], and more recent models use cancer-aware fusion or variational representations for incomplete inputs [8–9]. IntegrAO and joint variational autoencoders likewise demonstrate learning from partially observed molecular data [10–11]. SurvBoard highlights the value of evaluating survival methods under common protocols and of using patients with incomplete measurements [12]. These advances raise two connected questions: can a shared survival model learn effectively across different input combinations, and can predictions of the missing modalities add prognostic information to the measurements already available?

Molecular profiles pose a further representation problem: they contain heterogeneous, high-dimensional features without the spatial organization supplied by a tissue image. Table-to-image (T2I) methods encode feature relationships in a two-dimensional layout, allowing convolutional encoders to process neighboring features together [13–16]. Related approaches have used molecular image representations for survival prediction [17–18]. This provides a way to bring different molecular modalities into a common scheme for constructing inputs and pretraining. Pan-cancer learning also combines shared molecular structure with differences between cancer types [1,19]. A cancer-relative view can describe how a feature differs from the background of the same disease, alongside its standardized value and training-derived association with survival. We use these three views as aligned channels and evaluate risk ranking both across all patients and within cancer types.

Here we introduce OmniT2I, a unified table-to-image framework for survival prediction and cross-modal inference from naturally incomplete molecular and histopathology data (Fig. 1). Modality-specific encoders and availability-aware fusion enable a single model to use different input combinations; a target-excluded prediction path estimates withheld modalities from the observed ones. We evaluated patient-level out-of-fold predictions in 10,373 patients across 32 cancer types, comparing the shared model with separately trained survival methods across all 31 modality combinations. We also tested whether predicted modalities improve discrimination across 180 pairings of observed and withheld modality sets and evaluated frozen RNA-only transfer to seven external prognostic cohorts.

**Fig. 1.**
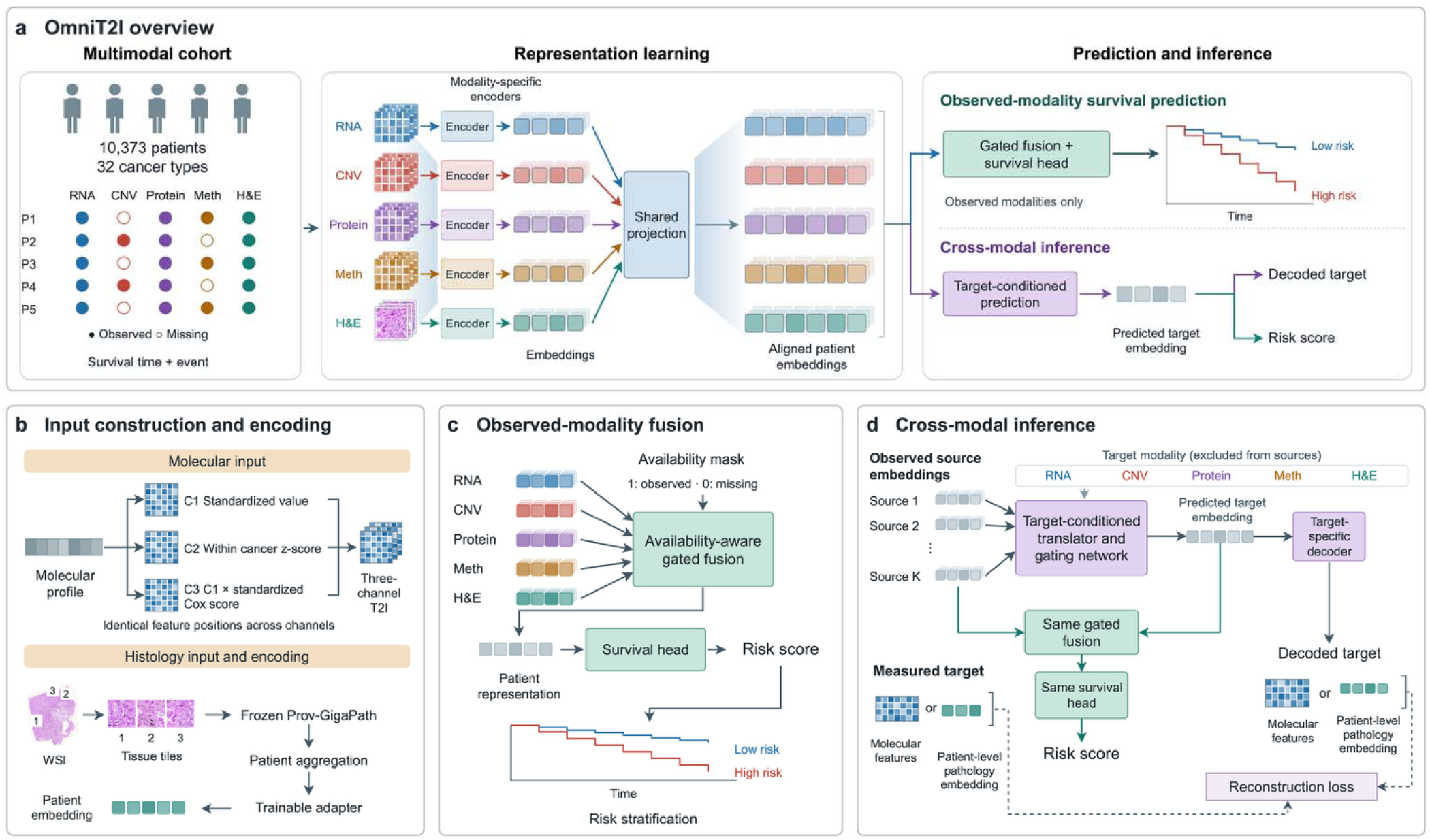
The OmniT2I framework. **a**, Overview of the naturally incomplete cohort, modality-specific encoding, shared projection and two inference tasks. Aligned patient embeddings support survival prediction from observed modalities and cross-modal inference. **b**, Molecular input construction and histopathology encoding. Three-channel molecular representations encode standardized values (C1), within-cancer standardized values (C2), and C1 weighted by standardized Cox score statistics (C3), with identical feature positions across channels. H&E tissue tiles are encoded using frozen Prov-GigaPath, aggregated at the patient level and processed by a trainable adapter. **c**, Availability-aware gated fusion combines observed embeddings, assigns missing modalities zero weight and produces a patient representation for the survival head. **d**, Target-conditioned translation predicts a target embedding from observed non-target modalities. The predicted embedding enters the same fusion and survival head alongside observed embeddings. A target-specific decoder reconstructs molecular features or a patient-level pathology embedding, supervised by the measured target during training. Survival curves are schematic.

## Results

### Overview of OmniT2I

OmniT2I represents each molecular profile as a three-channel T2I input (Fig. 1b). The channels encode standardized measurements (C1), within-cancer deviations (C2) and measurements weighted by standardized Cox score statistics estimated from training patients (C3). Each feature occupies the same position across channels. Masked reconstruction pretrains modality-specific encoders for RNA, CNV, protein and DNA methylation, whose readouts retain spatial positions. A frozen Prov-GigaPath encoder supplies histopathology features [20], which are aggregated at the patient level and processed by a trainable adapter. A shared projection maps the five modality embeddings into a common representation space (Fig. 1a).

The aligned embeddings support two prediction paths. For survival prediction, availability-aware gated fusion combines observed embeddings and assigns missing modalities zero weight; a survival head then produces a patient risk score (Fig. 1c). For cross-modal inference, target-specific mappings and source gating predict a selected modality embedding from the remaining observations. A target-specific decoder reconstructs molecular features or a patient-level histopathology representation (Fig. 1d). The measured target provides reconstruction supervision during training but is excluded from the predictor inputs. Predicted embeddings can also be combined with observed embeddings for survival prediction. During joint training, risk consistency encourages agreement between risk scores obtained with measured and predicted target embeddings.

### Multimodal coverage in the pan-cancer cohort

We characterized modality availability in the analysis cohort (Fig. 2). After sample selection and survival-data filtering, the cohort included 10,373 patients from 32 cancer types in The Cancer Genome Atlas (TCGA). Within this cohort, CNV inputs were available for 10,120 patients (97.6%), RNA for 9,550 (92.1%), and H&E for 9,539 (92.0%). DNA methylation and protein inputs were available for 8,183 (78.9%) and 7,262 (70.0%) patients, respectively. Only 5,605 patients (54.0%) had all five modalities. Another 3,439 patients (33.2%) had four, so 87.2% of the cohort contributed at least four modalities. RNA, CNV, DNA methylation, and H&E formed the most common incomplete set, with 2,200 patients (Supplementary Tables 1–3).

**Fig. 2.**
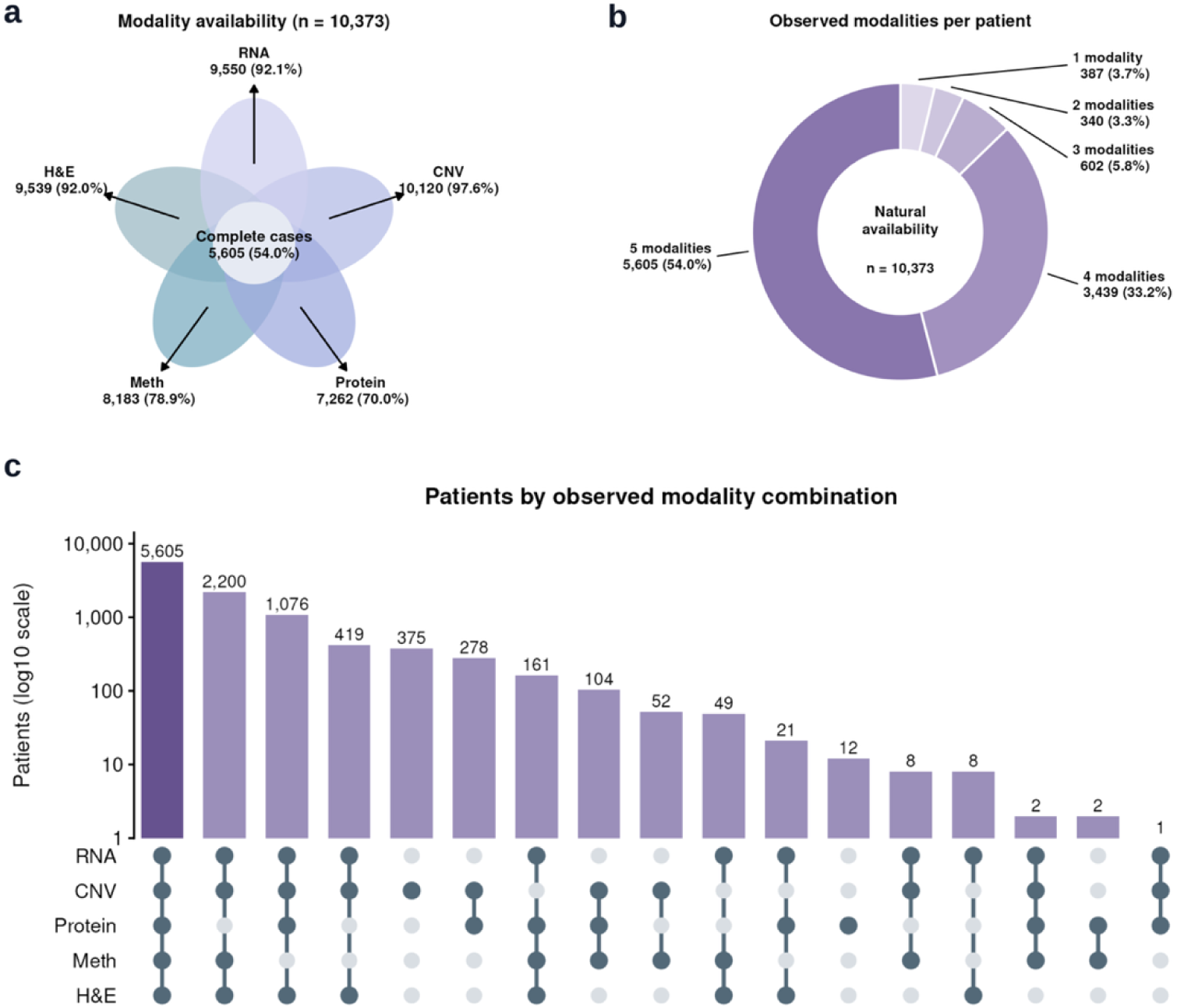
Naturally observed modality coverage in the TCGA cohort. **a**, Availability of RNA, copy-number variation (CNV), protein, DNA methylation (Meth) and histopathology (H&E) among 10,373 patients. Petals identify modalities; their area and overlap do not encode intersection sizes. The center reports 5,605 complete cases. **b**, Distribution of the number of measured modalities per patient. **c**, Counts of all naturally occurring availability combinations, with a log10 count axis above the aligned modality-membership matrix.

### OmniT2I predicts survival under natural modality availability

We first evaluated survival discrimination under natural modality availability using patient-level out-of-fold predictions from five outer folds and five random seeds (Fig. 3a). Across 10,373 patients and 3,258 deaths, the pooled concordance index (C-index) was 0.783 (95% confidence interval (CI), 0.776–0.790). In 5,605 complete cases with 1,546 deaths, it was 0.787 (0.778–0.797) (Supplementary Table 4).

**Fig. 3.**
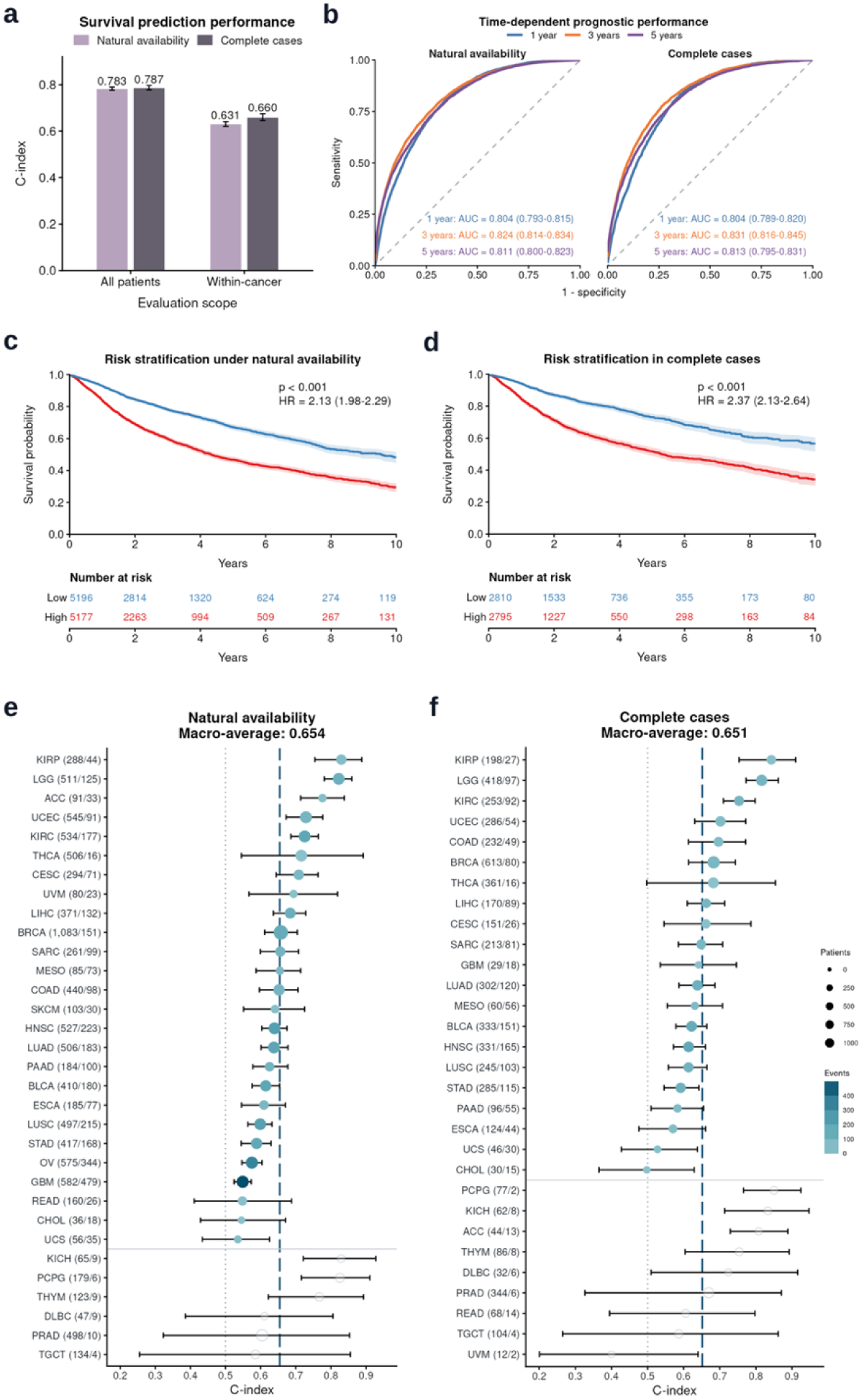
Prognostic discrimination and risk stratification under natural modality availability and in complete cases. **a**, Mean Harrell C-index across five seeds, pooled across cancers and restricted to within-cancer comparable pairs. Error bars are patient-bootstrap 95% percentile intervals. **b**, Pooled cumulative/dynamic receiver operating characteristic curves at 1, 3 and 5 years. Curves average outer-fold results within each seed and then five seeds; annotations give AUC and patient-bootstrap 95% intervals. Censoring distributions are estimated from outer-training patients. **c,d**, Kaplan–Meier curves with pointwise Greenwood log–log 95% bands and numbers at risk, for natural availability (10,373 patients; 3,258 deaths) and complete cases (5,605 patients; 1,546 deaths). Groups are defined by within-cancer median five-seed mean risk percentiles. Hazard ratios come from cancer-stratified Cox models and P values from cancer-stratified log-rank tests. **e,f**, Cancer-specific C-indices for all estimable cancers, ranked independently within eligible and low-count groups in each population. Point size represents patients and color encodes deaths for eligible cancers; semi-transparent gray points below the divider denote cancers with fewer than 25 patients or 15 deaths. Dashed lines show the unweighted mean among eligible cancers: 0.654 across 26 cancers in e and 0.651 across 21 cancers in f. There are 32 and 30 estimable cancers in e and f, respectively. All C-index and AUC intervals use 1,000 patient resamples, sharing sampled patients across seeds. Cancer type abbreviations are defined in Supplementary Table 2.

Concordance restricted to comparable patient pairs within the same cancer type was 0.631 (95% CI, 0.621–0.641) under natural availability and 0.660 (0.645–0.675) in complete cases. Within-cancer C-index reflects risk ranking among patients with the same cancer type. A cancer-type-only reference achieved pooled C-indices of 0.743 and 0.739, respectively (Supplementary Table 4); we therefore focus on within-cancer concordance below.

Pooled time-dependent areas under the receiver operating characteristic curve (AUCs) at 1, 3 and 5 years were 0.804, 0.824 and 0.811 under natural availability, and 0.804, 0.831 and 0.813 in complete cases (Fig. 3b; Supplementary Table 4).

Higher-risk groups had worse survival, with cancer-stratified Cox hazard ratios of 2.13 (95% CI, 1.98–2.29) under natural availability and 2.37 (2.13–2.64) in complete cases (Fig. 3c,d). Cancer-specific C-indices are shown for all estimable cancers: 32 under natural availability and 30 in complete cases (Fig. 3e,f). Among cancers with at least 25 patients and 15 deaths, unweighted mean C-indices were 0.654 for 26 cancers and 0.651 for 21 cancers, respectively. Cancer-specific time-dependent AUCs under natural availability and in complete cases are shown in Extended Data Fig. 1a and 1b, respectively (Supplementary Tables 5–7).

In an analysis of 6,386 patients from 21 cancer types (1,872 deaths), risk remained associated with overall survival (OS) after adjustment for age, sex and categorical stage in cancer-stratified Cox models. Fold-specific hazard ratios were 2.56–3.43 per unit of the five-seed mean standardized risk score, with all 95% CIs above 1. Sensitivity analyses retained the association (Supplementary Tables 8–9).

### One model supports multiple input combinations and competes with subset-specific specialists

We next compared the universal training strategy with nine survival methods fitted separately for all 31 nonempty modality combinations: DeepSurv, elastic-net Cox, lasso Cox, XGBoost Cox, random survival forest, survival gradient boosting, extra survival trees, CoxBoost and survival support vector machine. All 279 family-by-combination configurations used the same outer folds, seeds and training/validation assignments for patients included in both analyses. OmniT2I trained on naturally incomplete profiles, whereas specialists trained on complete cases; all were tested on the same 5,605 complete-case patients. Family-specific training and selection procedures are detailed in Methods (Supplementary Table 10).

OmniT2I achieved a higher mean within-cancer C-index in 226 of 279 family-by-combination comparisons (81.0%; Fig. 4a; Supplementary Table 11). Across the 31 combinations, OmniT2I’s mean within-cancer C-index exceeded that of five families by 0.014–0.052, with paired 95% CIs above zero. The corresponding differences for DeepSurv, elastic-net Cox, lasso Cox and XGBoost Cox were 0.001–0.005, with CIs including zero (Supplementary Table 12).

**Fig. 4.**
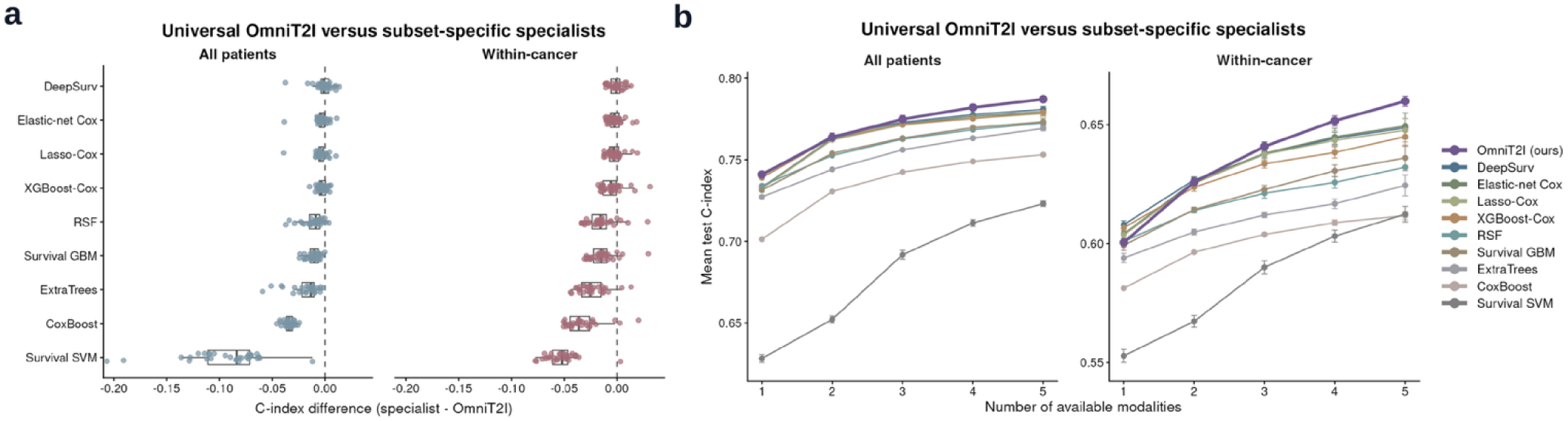
Universal and subset-specific survival models across measured modality combinations. **a**, Paired C-index differences for nine specialist families across all 31 nonempty combinations of five measured modalities, evaluated in the same 5,605 complete-case patients (1,546 deaths). Each point is one combination’s mean difference across five seeds; negative specialist-minus-OmniT2I values favor OmniT2I. Boxes show the median and interquartile range; whiskers extend to observations within 1.5 interquartile ranges. Facets show pooled and within-cancer concordance. **b**, Performance by number of available modalities. For each model and seed, C-indices are averaged equally over combinations of the specified size (5, 10, 10, 5 and 1 combinations); points and bars show the mean and one standard deviation across five seeds.

Mean within-cancer C increased from 0.600 with one modality to 0.626, 0.641, 0.652 and 0.660 with two, three, four and five modalities, respectively (Fig. 4b). OmniT2I had the highest combination-averaged performance among the evaluated models at three or more modalities; specialists remained competitive at smaller input sets. All individual combinations are reported in Extended Data Fig. 2 (Supplementary Tables 11–13). RNA had the highest mean within-cancer C-index among the five single modalities on these patients (0.633). Using RNA as one common reference, the five-modality model increased within-cancer C-index by 0.027 to 0.660 (Supplementary Tables 11–13).

### Predicted modalities retain prognostic information

We examined cross-modal prediction in complete cases, withholding each target while retaining its measurement for evaluation. Pairing each of five targets with the 15 nonempty subsets of the other modalities yielded 75 single-target routes. Reconstruction was assessed by feature-wise Pearson correlation across held-out patients and error relative to a training-derived cancer-mean reference. For H&E, the reconstruction target was the patient-level embedding (Supplementary Table 14).

Across-patient mean correlations were 0.739 for RNA, 0.681 for methylation, 0.610 for protein, 0.442 for CNV and 0.414 for H&E (Fig. 5a). After within-cancer centering, correlations were 0.228, 0.257, 0.187, 0.214 and 0.087, respectively. Across all test patients, reconstruction error was lower than that of the cancer-mean reference in 69 of 75 routes.

**Fig. 5.**
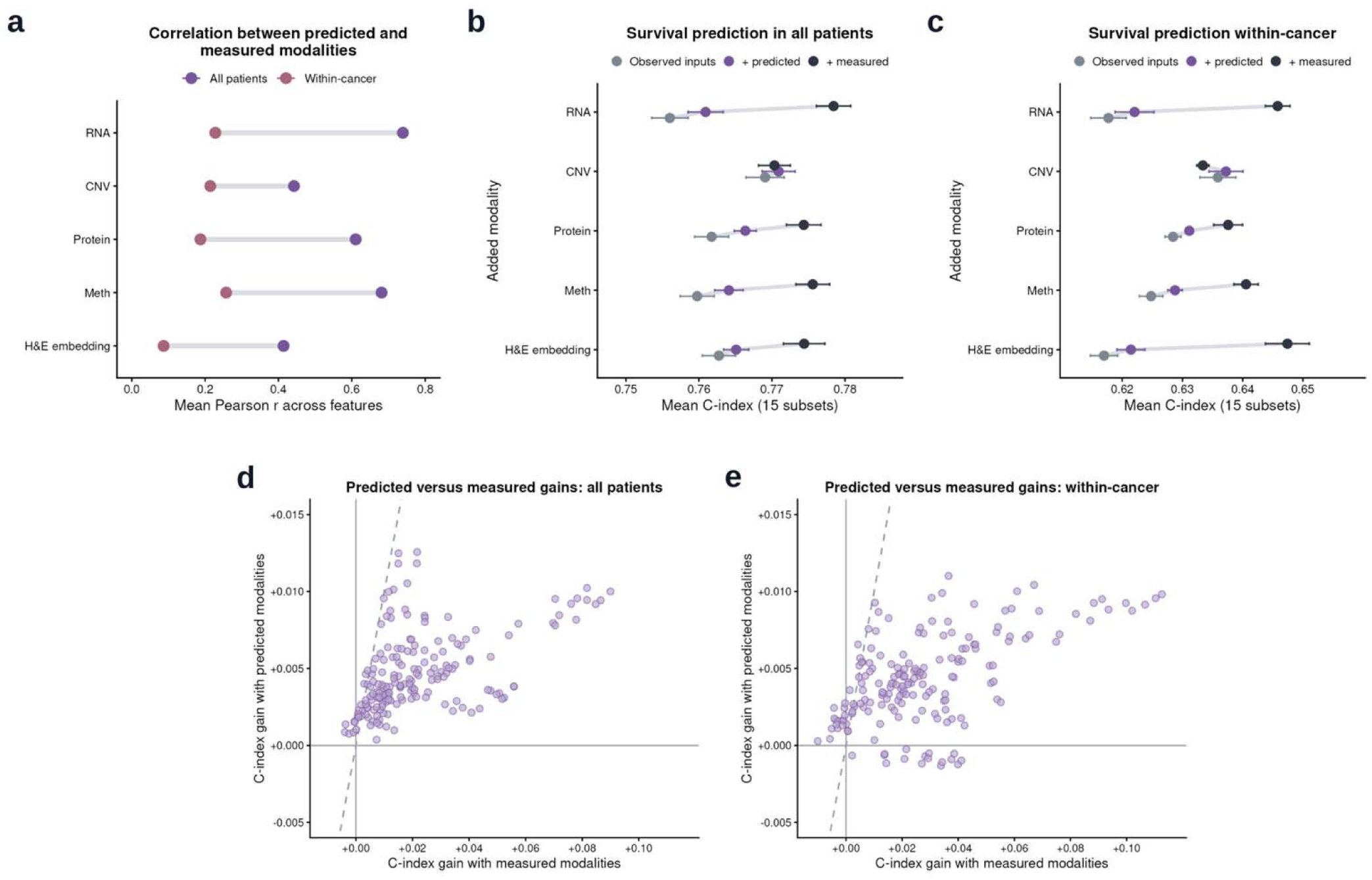
Reconstruction and survival utility of predicted modalities. **a**, Feature-wise Pearson correlation between predicted and measured targets, summarized across 15 nonempty source subsets excluding each target and averaged across five seeds. H&E reconstruction targets patient-level embeddings. **b,c**, C-indices across all patients (b) and within cancer types (c) for observed inputs alone, observed inputs plus one predicted target, and observed inputs plus the corresponding measured target. Observed inputs are measured modalities supplied to the model; predicted targets are inferred representations of withheld modalities, and measured targets are their representations from actual measurements. Points and bars show the mean and one standard deviation across five seeds after equally averaging 15 source subsets within each seed. **d,e**, Predicted-modality gains versus matched measured-modality gains across all 180 routes defined by source and target modality sets, shown across all patients (d) and within cancer types (e). Each point averages five paired seed-level differences. The measured reference contains the source–target union. Dashed diagonals indicate equal gains. Positive predicted gains occurred in all 180 routes across all patients and 164 routes within cancer types. All survival comparisons use the same 5,605 complete-case patients (1,546 deaths). “All patients” denotes pooled concordance. Averaged over all 180 routes, adding predicted modalities increased concordance across all patients by 0.0047 (95% CI, 0.0041– 0.0054) and within-cancer concordance by 0.0043 (0.0031–0.0054). Meth, DNA methylation.

Observed modalities are the measured inputs provided to the model and may include one or more modalities. Target modalities are withheld from these inputs. Predicted targets are model-generated representations of the withheld modalities; measured targets are their representations obtained from the actual measurements. For survival prediction, we compared observed inputs alone (A), observed inputs plus predicted targets (B), and observed inputs plus the corresponding measured targets, without refitting the model (Fig. 5b,c). Single-target predicted additions improved target-averaged within-cancer C by 0.001–0.004. Measured additions generally produced larger gains, but measured CNV reduced the target-averaged within-cancer C by 0.0025 while predicted CNV increased it by 0.0014 (Supplementary Tables 15–16).

Each route pairs a nonempty set of observed modalities with a nonempty set of withheld target modalities, with no modality shared between the two sets. The five modalities yield 180 such routes. Across these routes, mean B−A was +0.0047 across all patients (95% CI, 0.0041–0.0054) and +0.0043 within cancer (0.0031–0.0054). These gains corresponded to 22% and 15% of the mean gain from matched measured modalities, respectively. Gains were positive in all 180 routes across all patients and 164 within cancer (Fig. 5d,e). Predicted filling also exceeded cancer-mean filling in within-cancer C across all 180 routes. Of the 16 routes with negative within-cancer gains, 15 had H&E as the only observed source (Extended Data Fig. 3; Supplementary Table 16).

### Frozen RNA-only external evaluation across cancer cohorts

We evaluated the frozen RNA-only model in seven external prognostic cohorts comprising 3,901 patients and 607 deaths: six Clinical Proteomic Tumor Analysis Consortium (CPTAC) cohorts [21] and the Sweden Cancerome Analysis Network—Breast (SCAN-B; GSE96058) [22]. The external minus same-cancer TCGA RNA-only held-out test C-index (ΔC) ranged from −0.092 to +0.079, with a median of −0.003; three estimates were higher and four lower (Fig. 6a; Supplementary Table 17). SCAN-B, the largest cohort with 3,273 patients and 336 deaths, yielded C = 0.637 (95% CI, 0.606–0.668), compared with 0.639 in TCGA BRCA (ΔC = −0.003). The largest negative difference occurred in CPTAC HNSCC (ΔC = −0.092), a cohort negative for human papillomavirus (HPV) [23].

**Fig. 6.**
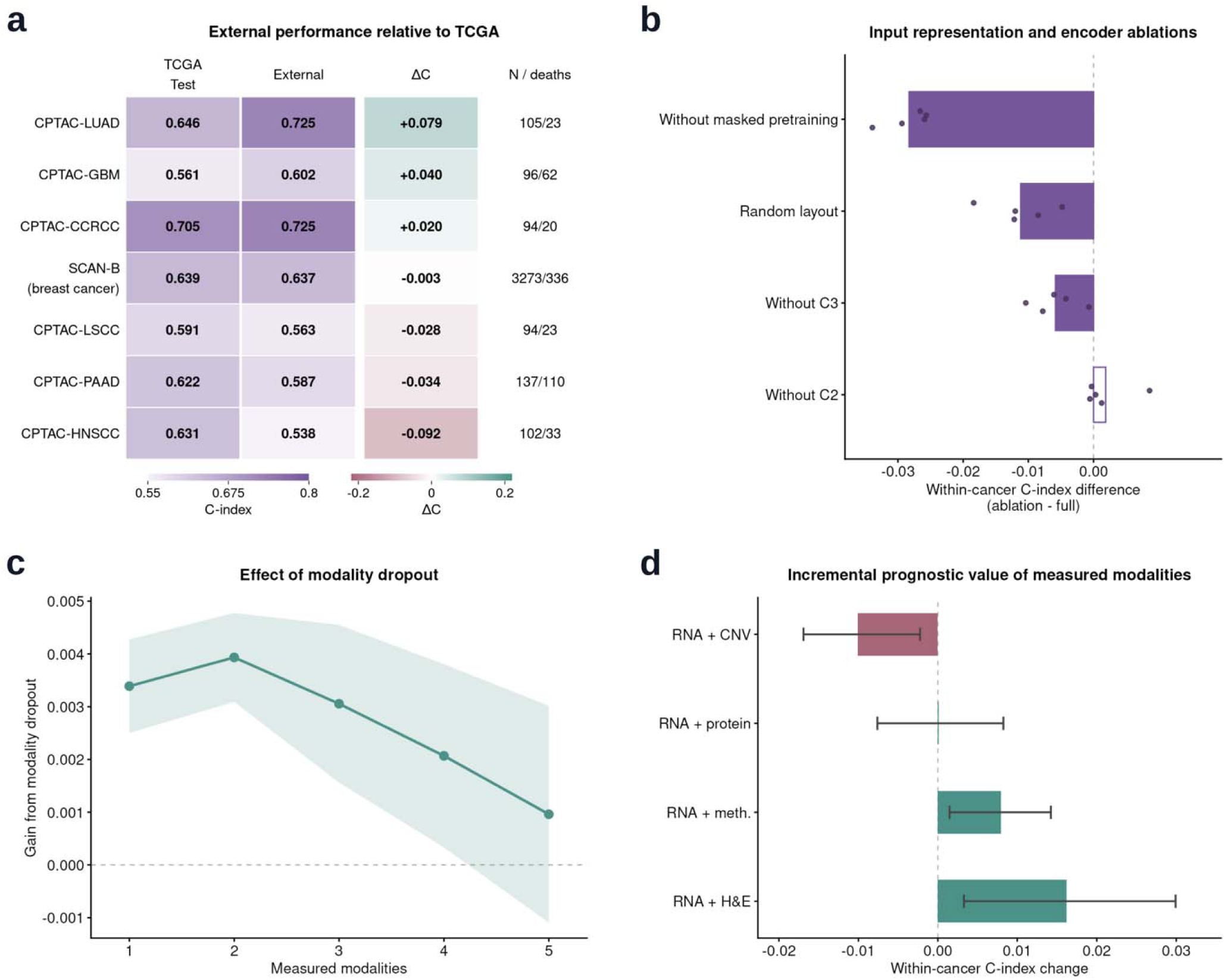
External evaluation, component ablations and modality contributions. **a**, Frozen RNA-only performance in seven external prognostic cohorts. Columns show same-cancer TCGA held-out test C-index, external C-index and ΔC, defined as external minus TCGA test C-index. Right labels give external patients/deaths. TCGA references use five-seed out-of-fold predictions; external risks average 25 frozen models. Pointwise 95% intervals are reported in Supplementary Table 17. **b**, Input representation and encoder ablations under natural availability (10,373 patients; 3,258 deaths). Bars show five-seed mean within-cancer differences (ablation minus full model); dots show paired seed differences. Purple fill denotes a negative mean with decreases in at least four seeds; other bars are open. **c**, Gain from modality dropout in 5,605 complete cases (1,546 deaths). Within each seed, differences are averaged over 5, 10, 10, 5 and 1 combinations for one to five measured modalities. The line shows the five-seed mean; shading shows ± one between-seed s.d. **d**, Within-cancer C-index change after adding one measured modality to RNA in the same complete cases. Bars show five-seed means; error bars show 95% paired patient-bootstrap intervals from 2,000 resamples. Teal and rose denote non-negative and negative changes. RNA-only C-index is 0.6334.

### Contributions of pretraining, input representation and modality availability

Removing masked pretraining lowered natural-availability within-cancer C by 0.028, removing C3 by 0.006, and replacing the correlation-based layout with a fixed random layout by 0.011 (Fig. 6b). Removing C2 or risk consistency, or replacing gated fusion with averaging, had small effects on this endpoint. Under controlled missingness, removing modality dropout lowered within-cancer C in all 31 measured subsets, by 0.0032 on average across the 31 subsets, with each subset weighted equally after averaging its difference across five seeds (Fig. 6c); removing pretraining lowered all 31, by 0.0177 on average. C2 increased pooled C in 28 of 31 measured subsets, by 0.0046 on average, while within-cancer C was lower by 0.0020. The gated-fusion model had higher mean within-cancer C than the mean-fusion model with one or two observed modalities, by 0.0018 and 0.0008, respectively; mean fusion performed slightly better with three or more. Risk consistency increased mean predicted-modality gain from 0.0040 to 0.0043 within cancer and from 0.0038 to 0.0047 pooled (Supplementary Table 18).

The contribution of an additional measurement depended on the modality. In the same complete-case patients, adding H&E to RNA increased within-cancer C by 0.016 (95% CI, 0.003–0.030), and methylation increased it by 0.008 (0.001–0.014). Protein produced no clear increment (+0.0001; −0.008 to +0.008), whereas CNV decreased C by 0.010 (−0.017 to −0.002; Fig. 6d). Results for all 75 single-modality additions are provided in Supplementary Table 19.

## Discussion

OmniT2I brings incomplete molecular profiles and histopathology into a shared framework for prognostic learning. One model accommodated all 31 nonempty input combinations and achieved higher within-cancer C-indices in 226 of 279 comparisons with nine separately trained survival methods. Predicted modalities added prognostic information in 164 of 180 observed–withheld modality pairings, and the frozen RNA-only model was evaluated in seven external cohorts.

Previous T2I methods, including DeepInsight, IGTD and OmicsFootPrint, organize tabular features into images for convolutional learning [13,15–16]. Molecular image representations have also been investigated for survival prediction [17–18]. OmniT2I uses modality-specific encoders for four molecular modalities and integrates their representations with a pathology representation from Prov-GigaPath [20]. Correlation-based feature placement supplies a spatial arrangement, while C1–C3 provide aligned views of standardized measurements, cancer-relative deviations and survival-associated weighting. Masked pretraining then applies a common learning task across the molecular encoders. Removing pretraining, replacing the correlated layout with a random layout or removing C3 reduced natural-availability within-cancer C by 0.028, 0.011 and 0.006, respectively. These findings support the organization and pretraining of molecular inputs as components of the framework. C2 provides disease-relative context, although its contribution was mainly evident in pooled discrimination.

Our shared-model results complement studies of multimodal integration, including PORPOISE, MCAT, SurvPath and PROGPATH [3,5–6,24], and benchmark evidence that incomplete profiles can contribute to survival-model training [7,12]. The nine-method comparison within our study supplies the direct performance evidence under shared evaluation conditions. Its findings reflect a training strategy that includes patients with incomplete profiles, alongside shared model parameters. External RNA-only evaluation adds a separate test of transfer: ΔC relative to same-cancer TCGA test performance had a median of −0.003 and ranged from −0.092 to +0.079. SCAN-B provided the largest cohort and a ΔC of −0.003 [22]. The largest decrease occurred in the HPV-negative CPTAC HNSCC cohort [23]; differences in population composition motivate subgroup evaluation.

Cross-modal inference extends the analysis from integrating measured modalities to predicting information that is unavailable for a patient. Related work has explored cross-modal translation and alignment for survival analysis [25], masked prediction of molecular and pathology representations [26], and molecular imputation or translation in other settings [11,27]. OmniT2I evaluates both reconstruction and the survival benefit of adding predicted representations to observed inputs. Reconstruction error was lower than the cancer-mean reference in 69 of 75 routes across all test patients, while the mean within-cancer survival gain was 0.0043, corresponding to 15% of the gain from matched measured modalities. These measures answer different questions about the inferred representations. H&E supplied the largest measured addition to RNA, yet it was the sole observed source in 15 of the 16 routes with negative within-cancer augmentation gains. Predicting other modalities from histology and using histology alongside molecular measurements therefore place different demands on the learned representation. This distinction also complements evidence that the value of multimodal fusion depends on how its component predictions are combined [28].

Several limitations guide interpretation and further evaluation. TCGA and the external cohorts were assembled retrospectively, and complete-case comparisons may not represent patients with naturally missing assays. The specialist comparison combines differences in model design and the training population. External data differed in assay processing and follow-up, and several cohorts had few deaths. The external analyses tested RNA-only prognosis; comparable external evaluation of multimodal integration and cross-modal inference remains necessary. Incremental prediction beyond established clinical models and calibration of absolute survival probabilities also remain to be assessed.

Future work can extend the representation to clinical variables and radiographic images, bringing additional scales of patient information into the shared model. Larger paired datasets could support richer cross-modal prediction and evaluation across patient subgroups, while longitudinal measurements could connect changes in molecular and tissue states to prognosis. Prospective studies will be needed to establish how these predictions contribute to clinical workflows.

OmniT2I combines table-to-image molecular representations, histopathology and cross-modal inference in one survival model for incomplete multimodal cancer data. Its shared training strategy accommodates diverse input combinations, and its measured- and predicted-modality evaluations quantify how additional information changes prognostic discrimination.

## Methods

### Study design and data sources

This retrospective computational study used public, de-identified data from TCGA Pan-Cancer Atlas [1] and external cancer cohorts, including CPTAC [21]. TCGA overall survival outcomes and cancer-type labels were obtained from the TCGA Pan-Cancer Clinical Data Resource [29]. Molecular matrices were downloaded from the University of California Santa Cruz (UCSC) Xena platform [30]. RNA inputs were batch-normalized abundances. Copy number inputs were continuous gene-level GISTIC2 values [31]. Protein and DNA methylation inputs were normalized reverse phase protein array (RPPA) abundances and 450K β-values, respectively. Open-access H&E whole-slide images (WSIs) and case metadata were obtained from the National Cancer Institute Genomic Data Commons (GDC) [32]. External sources and evaluation procedures are described below and in Supplementary Table 20.

### Cohort assembly, sample types and survival endpoints

Molecular modalities were matched at the patient level using one Primary Tumor profile per modality per patient. For H&E, all open, released Primary Tumor diagnostic slides were retained. The existing Primary Tumor tissue-slide selection was used for patients without an available diagnostic slide. H&E collection was conditional on RNA availability.

Overall survival was defined by the TCGA Clinical Data Resource OS indicator and positive OS.time in days. Records with missing or non-positive follow-up, or event values outside 0 or 1, were excluded. Cancer type was defined by the TCGA study abbreviation. The union cohort comprised patients with valid endpoints and at least one observed modality; complete cases had all five modalities. Cohort sizes, event counts and availability patterns are provided in Supplementary Tables 1–3.

### Patient-level outer cross-validation

A patient-level fivefold partition was stratified by cancer type and event status and shared across modalities (Supplementary Table 22).

Each outer training set was divided into 80% core training and 20% inner validation patients, stratified by cancer type and event status (Supplementary Table 22). The same patient-level assignments were reused across all training stages. Feature selection, imputation, C1/C2 normalization and spatial ordering were fitted using outer-training patients, including inner validation covariates. Outcome-derived C3 weights were fitted using only core patients with the measured modality.

Trainable model parameters and cancer-specific decoder means were fitted using core training patients. Inner validation outcomes selected survival checkpoints and contributed to the joint-training survival constraint. Outer test risk scores were standardized using only the mean and population standard deviation of inner validation risk scores. Five random seeds varied inner partitioning, initialization and training stochasticity. The same outer folds were used for model development, architecture comparisons and reported evaluation.

### Three-channel molecular input construction

Each molecular modality was represented by three channels with identical feature positions. Channel 1 contained standardized assay values. Channel 2 contained feature-wise z-scores of Channel 1 within each cancer type, with reference statistics fitted using outer training patients. Channel 3 multiplied Channel 1 by a feature-wise Cox partial-likelihood score evaluated at a zero coefficient, computed only from measured core patients. For each event, this score subtracts the feature mean among patients still at risk from the event patient’s feature value, then sums across events. Tied events share the full risk set, including patients censored at that time, following the

Breslow convention. Scores were standardized across features and clipped to [−3, 3]; they were not fitted Cox regression coefficients. The same weights were applied to core, validation and outer test patients. Features were ordered by average-linkage clustering of one minus their outer-training correlation. Ordered features were placed along a serpentine path for all four molecular modalities, with identical feature positions across the three channels. RNA, CNV and DNA methylation used 3 × 45 × 45 inputs; protein used 3 × 17 × 17 inputs. Unoccupied positions were zero-padded.

### RNA expression preprocessing and T2I construction

RNA values were supplied as batch-normalized log2(normalized abundance + 1). Duplicate feature identifiers were averaged, and identifiers with missing values across eligible RNA samples were removed. Feature identifiers were retained as supplied by Xena.

RNA expression was normalized against a fold-specific invariant panel of low-variance features spanning three expression bins. Let *x_ig_* denote the batch-normalized expression of gene *g* in patient *i* and let *I_f_* denote the panel selected from outer training fold *f* The sample factor was

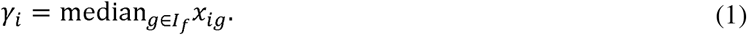

The corrected expression was

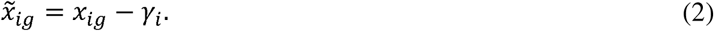

Channel 1 was then defined as

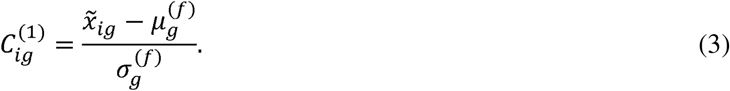

Here,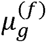 and 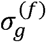 are the outer training mean and population standard deviation of corrected expression; values of 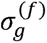 below 10^-6^were set to one. The 2,000 genes with the highest variance after invariant normalization were retained. For an unseen cancer type, RNA Channel 2 used global training statistics.

### CNV preprocessing and T2I construction

The CNV branch used continuous gene-level GISTIC2 values from eligible Primary Tumor patients. Within each outer training fold, the 2,000 highest-variance features were selected and standardized using their training means and population standard deviations.

### Protein abundance preprocessing and T2I construction

The protein branch used normalized RPPA measurements of total proteins and phosphoproteins. Analytes were filtered by missingness and variance within each outer training fold. Within available protein profiles, missing analyte values were filled with the corresponding outer-training analyte medians to provide complete encoder inputs. Retained values were then standardized using outer-training means and standard deviations. Entirely unmeasured modalities were handled by the availability mask. The same 217 analytes passed quality control in all five folds.

### DNA methylation preprocessing and T2I construction

450K probes were mapped to protein-coding promoters using GENCODE version 19 on hg19. Promoters extended from 1,500 base pairs upstream to 500 base pairs downstream of each transcript start site, accounting for strand orientation. Probe midpoints were intersected with promoters, and probes assigned to multiple genes were excluded. Genes required at least three unambiguous promoter probes. Technical columns for the same patient were combined by their observed mean. Gene-level β-values were obtained by averaging observed promoter probes.

Gene-level β-values were clipped to [10^-5^,1-10^-5^] and transformed to M-values as *M =* log_2_[*β* / (1 – β)]. Chromosome Y genes were excluded. Within each outer training fold, genes were filtered by missingness and variance. Missing gene-level M-values within available methylation profiles were filled with the corresponding training-fold gene medians to provide complete encoder inputs. The 2,000 highest-variance genes were retained and standardized to form Channel 1.

### Histopathology representation

H&E tissue regions were sampled from the selected slides and filtered by tissue fraction before encoding. Retained patches were resized, center-cropped and normalized with ImageNet channel statistics.

The frozen Prov-GigaPath tile encoder generated a representation for each retained patch [20]. Patch representations were averaged within slides, then averaged with equal slide weights to obtain a 1,536-dimensional patient representation. Patient vectors were standardized using outer training H&E statistics and clipped before entering the trainable adapter. H&E used this patient representation independently of molecular T2I construction. Preprocessing specifications, channel definitions and input dimensions are provided in Supplementary Table 21.

### Separate encoders for each modality and a shared latent space

Each molecular modality used an independent encoder with three 3 × 3 convolutional blocks, batch normalization and rectified linear activation. The first two blocks included max pooling. A 1 × 1 convolution reduced the channels of the final spatial feature map, followed by Gaussian error linear unit (GELU) activation. All spatial positions were retained, flattened and mapped by a position-specific linear readout to a 96-dimensional embedding.

In stage A, molecular encoders underwent masked reconstruction pretraining using only core patients with the relevant modality. For each patient, 15% of measured feature positions were randomly masked across all three channels. A temporary linear decoder reconstructed the masked values using smooth L1 loss, excluding padding and unmasked features. Pretraining was followed by supervised unimodal Cox fitting on the same core patients, with checkpoint selection on the same inner validation patients. These supervised encoders initialized stage B multimodal survival training; the temporary reconstruction decoder was not retained.

A trainable two-layer adapter mapped the frozen 1,536-dimensional H&E patient representation to a 96-dimensional embedding. Layer normalization, GELU activation and dropout were applied within the adapter.

A projection head with shared parameters mapped each modality embedding into a common 96-dimensional latent space. A separate linear auxiliary risk head operated on each projected modality representation. The fused survival head mapped the combined patient representation to a scalar risk score.

### Gated fusion that accommodates missing modalities

Let *M* denote the set of five modalities. Let *z_im_* ∈ ℝ^96^ denote the projected representation of patient *i* from modality *m*, and let *a_im_* ∈{0, 1} indicate whether that modality was observed. A separate linear gate for each modality produced logit *q_im_* = *g_m_*(*z_im_*)The normalized weight was

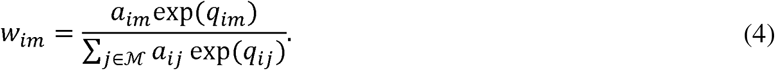

The fused representation was

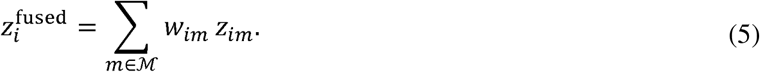

Unavailable modalities received zero fusion weight, and weights for the retained modalities summed to one. During training, whole-modality dropout modified the availability mask while retaining at least one observed modality per patient. This operated independently of feature masking during encoder pretraining.

### Translation across modalities and decoders for measured features

The translation module contained 20 directed source-to-target routes, each combining a low-rank linear skip with a nonlinear residual branch. For target *t*, the target modality was excluded from the source mask. Target-specific route gates were normalized across available non-target sources. Their weighted outputs formed a predicted 96-dimensional target embedding.

A target-specific decoder mapped the predicted embedding to the selected molecular feature profile or the 1,536-dimensional H&E patient representation. Targets were standardized using means and standard deviations estimated from outer-training patients with the corresponding measured modality in each fold. Decoders predicted residuals around cancer-specific target means estimated from core training patients. Reconstruction loss was evaluated only when the target was measured.

Cross-modal reconstruction provided auxiliary supervision during training. At primary survival inference, fusion used only the patient’s observed modalities.

### Survival and multitask objectives

The survival objective combined the negative Cox partial log-likelihood of the fused risk, the mean Cox loss of available unimodal auxiliary heads and an alignment loss:

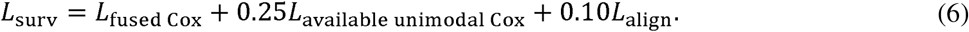

The auxiliary Cox term for a modality was computed only among patients for whom that modality was observed. The alignment term was the mean squared distance between shared-projection embeddings, averaged across modality pairs observed in the same patient.

For training across modalities, the observed target embedding was detached and used only as supervision. Reconstruction of both the target latent and measured features combined smooth L1 loss with cosine distance penalties weighted by 0.10 and 0.05, respectively. A consistency term used smooth L1 loss to match the risks obtained when the true target latent or its predicted latent was inserted into the same deterministic fusion risk map. These three losses were averaged with equal weights across eligible targets. We defined

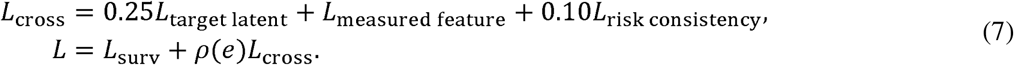

The ramp coefficient was

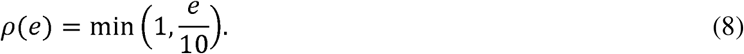

Here, e denotes the joint-training epoch index. During decoder warm-up, the survival backbone was fixed and *L*_cross_ was minimized without ramping. During joint training, the model minimized the complete objective above, and ρ(*e*)increased linearly over the first ten joint training epochs. This construction allowed reconstruction gradients to reach the trainable parts of the molecular encoders while preventing an identity route from the target itself.

### Optimization and implementation

Training comprised masked pretraining and supervised unimodal fitting in stage A, followed by multimodal survival training in stage B. Stage C first trained the translator and decoders with the survival backbone frozen, then jointly optimized them with selected survival-model components; earlier molecular layers, spatial readouts, batch-normalization running statistics and the Prov-GigaPath backbone remained frozen. Architecture, optimizer settings, training budgets and checkpoint selection are detailed in Supplementary Table 22.

OmniT2I was implemented in Python 3.10 using PyTorch 2.4.1. The final model and component ablations used five seeds and five outer folds.

### Component ablations

Seven individual ablations removed masked pretraining, C2, C3, survival modality dropout or risk consistency; replaced gated fusion by averaging over available modalities; or randomized molecular layouts. A joint averaging/no-risk-consistency variant was also evaluated. Removing a channel set it to zero throughout A–C and excluded it from masked reconstruction loss while retaining network dimensions. Random layouts used one fixed permutation per modality and outer fold, shared across seeds; all channels moved together and padding was unchanged. Each ablation retrained all stages affected by the change (Supplementary Table 22). Translation source dropout was retained. All other inputs, partitions, supervised budgets and checkpoint rules were held fixed, and all variants were evaluated on natural availability, 31 measured subsets and 180 predicted-modality routes.

### Evaluation and statistical analysis

Performance was evaluated from patient-level out-of-fold predictions under natural modality availability and in complete cases. Predictions from five outer folds were combined separately for each of five seeds. Before pooling, risk scores were standardized against the corresponding inner validation predictions, separately for the two evaluation conditions.

The primary discrimination metric was Harrell’s C-index [33]. Evaluation across all patients used all eligible comparable patient pairs. Evaluation within cancer restricted comparable pairs to patients with the same cancer type. Time-dependent prognostic discrimination at 1, 3 and 5 years was quantified using cumulative dynamic AUC with inverse probability of censoring weights [34], implemented in scikit-survival 0.25.0 [35]. The censoring distribution was estimated from the corresponding outer training fold. AUC estimates for each fold were weighted by the number of eligible test patients and then averaged across folds and random seeds.

For the cancer-type-only reference, we divided the total number of deaths by the total follow-up time within each cancer type in each outer training fold. Each held-out patient was assigned the resulting value for their cancer type as a risk score. We combined predictions across the five outer test folds and calculated pooled Harrell’s C-index under natural availability and in complete cases. The complete-case analysis restricted only the evaluation population; both analyses used reference scores estimated from all outer-training patients (Supplementary Table 4).

Confidence intervals for the principal C-index and AUC estimates were obtained from 1,000 patient bootstrap resamples. Patients were sampled with replacement within cancer type, and the same sample was applied to predictions from all five seeds. Metrics were calculated for each seed and averaged before two-sided 95% percentile intervals were derived. Models were not retrained. Cancer-specific intervals resampled patients within each cancer. These intervals condition on fitted predictions and do not include refitting, tuning or development-stage model-selection uncertainty.

For risk stratification, out-of-fold risk scores were converted to percentile ranks separately by cancer and seed, using (average rank − 0.5)/n, where n is the number of evaluated patients in the corresponding cancer type. Patient ranks were averaged across seeds; scores above the cancer-specific median defined higher risk, and scores at or below it defined lower risk. Kaplan–Meier curves used pointwise Greenwood log–log 95% intervals, and a two-sided cancer-stratified log-rank test compared groups. Hazard ratios and Wald intervals came from an unpenalized Cox model stratified by cancer, using Efron ties, fitted with lifelines 0.30.0.

### Clinically adjusted prognostic associations

Clinically adjusted prognostic associations were assessed among patients with recorded age, sex and pathologic stage I–IV. Each patient contributed the mean of five out-of-fold risk scores standardized using inner validation predictions under natural modality availability. Unpenalized Cox models were fitted separately in each outer test fold, stratified by cancer and adjusted for age per decade, sex and categorical stage. Hazard ratios refer to a one-unit increase in the mean score, not one standard deviation of the ensemble. Fold-specific 95% intervals and two-sided nominal Wald P values are reported in Supplementary Table 8. Rank-time proportional-hazards diagnostics (Supplementary Table 9) and three retained sensitivity specifications were examined: a cubic age spline, cancer-by-stage strata, and cancer-by-stage strata with an age effect changing after one year. OmniT2I was not refitted. These analyses assess adjusted association, not added predictive utility.

### Universal versus subset-specific survival models

The universal OmniT2I model was trained with modality masking in the naturally incomplete cohort of 10,373 patients. One universal model was fitted for each cross-validation fold and random seed and applied to all 31 nonempty combinations of the five modalities. Comparator models were trained separately for each combination using the 5,605 complete cases. Every model was evaluated on the same 5,605 complete cases for this benchmark. The benchmark included DeepSurv [36], elastic-net Cox, lasso Cox, XGBoost Cox, random survival forest, survival gradient boosting machine, ExtraTrees, CoxBoost and survival support vector machine. DeepSurv used PyTorch 2.4.1; XGBoost Cox used XGBoost 3.2.0 [37]; the remaining families used scikit-survival 0.25.0 [35]. CoxBoost was implemented as componentwise gradient boosting in scikit-survival.

Shared patients retained identical outer folds, endpoints, seeds and saved core/validation assignments. For specialists, these assignments were intersected with complete-case membership, rather than constructing a separate split after restricting patients. Specialists received fold-specific conventional feature vectors for their designated modalities. Preprocessing used outer training data, with additional feature standardization on core training patients for elastic-net Cox, lasso Cox, CoxBoost and survival SVM. Model fitting used complete-case core training patients, and risk standardization used inner validation predictions. Cox penalty strengths were selected along a regularization path using validation C-index. Neural checkpoints and XGBoost iteration counts used validation-based early stopping; other family-specific settings were fixed. Family-specific training and selection settings are provided in Supplementary Table 10.

Primary comparisons used each of the nine survival methods, not the family with the best observed test score for each subset. Differences were calculated within modality combination and seed. Figure 4 reports specialist minus OmniT2I; negative differences favor OmniT2I. Family-level effects average equally over 31 combinations and five seeds. Modality-count summaries average equally over combinations of the same size within each seed. Seed standard deviations describe variation across fitted runs and are not patient-sampling confidence intervals. Subset-specific performance, paired confidence intervals and modality-count summaries are provided in Supplementary Tables 11–13.

### Paired uncertainty and conditional measured-modality gains

Paired 95% percentile intervals used 2,000 patient bootstrap resamples within cancer for each fixed-family comparison averaged over 31 subsets and for the B−A effect averaged over 180 routes. Seed-specific C-indices were calculated before averaging; risk scores were not averaged for these contrasts. Within-cancer C pooled comparable pairs within cancers rather than averaging cancer-specific C (Supplementary Tables 12, 16, 19).

For measured modalities, all 75 transitions from a nonempty source set S to S plus one absent modality m were summarized as C(S+m)−C(S), using the same 5,605 complete cases. Each transition reports the five-seed mean and number of positive seeds. Four comparisons of RNA alone with RNA plus CNV, protein, DNA methylation or H&E received the same paired intervals from 2,000 resamples. All 75 transitions are provided in Supplementary Table 19; the intervals for these four additions to RNA accompany Fig. 6d (Supplementary Table 19).

### Cross-modal prediction and prognostic gain

Cross-modal prediction was evaluated in complete cases, where every target was measured. Each target was predicted from every nonempty subset of the remaining four modalities, giving 5 × (2^4^ – 1) = 75 single-target routes. The target was excluded from the inputs in every route. Predicted targets are model estimates generated from the observed inputs; measured targets are the corresponding patient measurements used to evaluate reconstruction and the survival benefit of adding the actual modalities.

Prediction accuracy was quantified by calculating Pearson correlation between predicted and measured values across outer test patients for each target feature and then averaging across features. Fold estimates were weighted by the number of target test patients. Within-cancer correlations were calculated after centering predicted and measured values within each outer test fold and cancer; groups with fewer than two patients were excluded. For each target and seed, correlations were averaged equally across its 15 source subsets. Figure summaries show means and sample standard deviations across five seeds. Stored cancer-specific target means estimated from core training patients provided a separate reconstruction reference. H&E targets were patient embeddings, not image pixels. Route-level reconstruction results are provided in Supplementary Table 14.

Prognostic gain was assessed using observed inputs alone (A), observed inputs plus predicted target embeddings (B), and observed inputs plus the corresponding measured target embeddings. The matched measured reference was obtained from the 31-subset evaluation for the exact source–target union. C-index differences were calculated relative to sources alone, keeping trained encoders, projection, fusion and survival head fixed. Predicted target latents entered fusion directly; decoded features were not re-encoded. Each target was inferred independently from the original measured sources, never from another predicted target.

Multi-target analyses included all 180 ordered routes with nonempty, disjoint source and target sets. Each route was summarized across five seeds. The primary augmentation contrast was B−A, equally averaged over all routes. The descriptive ratio of mean gains divided the equally weighted mean predicted-target gain by the corresponding mean gain from matched measured targets across all 180 routes, rather than averaging individual route ratios. Comparisons with cancer-mean filling were auxiliary, distinct from B−A. Route-level survival results and average gains with paired confidence intervals are provided in Supplementary Tables 15–16.

### Extended Data analyses

Cancer-specific time-dependent AUC was estimated independently at 1, 3 and 5 years. A cancer and time horizon was displayed when it contained at least 10 cases, at least 10 controls and evaluable estimates from at least three outer folds. Cancer types were ordered independently at each time horizon according to AUC. Confidence intervals were calculated using 1,000 patient bootstrap resamples within each cancer type.

Fig. 3e,f includes all cancers with estimable C-indices: 32 under natural availability and 30 in complete cases. Within each population, cancers meeting the minimum of 25 patients and 15 events are ranked by descending C-index, followed below a separator by other estimable cancers shown as semi-transparent points. Macro-averages include only the 26 eligible cancers under natural availability and 21 in complete cases; their values are 0.654 and 0.651, respectively. Complete-case OV and SKCM have no patients and are non-estimable, so they are omitted. Cancer-specific intervals used 1,000 patient bootstrap resamples within cancer. The benchmark heatmap reports absolute C-index for all models and modality combinations, whereas cross-modal heatmaps report B−A. Values are means across five seeds; routes and subsets share patients. Cancer-specific discrimination, AUC reporting status, subset-specific performance and predicted-modality gains are reported in Supplementary Tables 6–7, 11 and 15–16.

### Frozen RNA-only external evaluation

External evaluation included prognostic cohorts with baseline tumor RNA, evaluable OS in at least 25 patients and at least 15 deaths. Six CPTAC cohorts (LUAD, CCRCC, GBM, PAAD, LSCC and HNSCC) and SCAN-B met these criteria. RNA profiles and outcomes were matched by patient identifier; invalid or non-positive survival times were excluded. Results, cohort-specific OS definitions and public access routes are provided in Supplementary Tables 17, 20.

Expression adapters retained deposited logged matrices without an additional log transformation, or converted linear abundances once to the required log scale. For SCAN-B, fragments per kilobase of transcript per million mapped reads (FPKM) values were recovered from the original log2(FPKM + 0.1) release, scaled to transcripts per million (TPM), and transformed to log2(TPM + 1); explicitly labeled replicate columns were excluded (Supplementary Table 20). Exact symbols or unambiguous one-to-one renamings were mapped to each checkpoint’s fixed feature positions. For every external sample, the median expression of measured genes in the saved invariant panel was subtracted from log-scale expression. Genes missing from an external RNA matrix were filled with their saved TCGA training means on this corrected scale to retain the checkpoint’s fixed input panel. The inputs were then processed using the stored global C1 standardization, the corresponding TCGA cancer C2 reference, corrected core-derived TCGA OS C3 weights and the saved layout. No external outcomes or cohort-level normalization parameters were fitted. Each frozen checkpoint’s risk was standardized by its saved TCGA natural-risk mean and standard deviation; 25 fold-by-seed risks were averaged per external patient. Pointwise 95% C-index intervals used 2,000 patient-bootstrap draws conditional on these predictions; unevaluable draws were omitted.

Same-cancer TCGA references supplied RNA alone to the model, using patients with measured RNA and CNV; counts are reported in Supplementary Table 17. TCGA test C-indices used each patient’s mean risk across five seed-specific models for which that patient was in the held-out outer fold. External predictions averaged all 25 fold-by-seed models. ΔC was the external C-index minus the same-cancer TCGA test C-index.

CPTAC follow-up was generally diagnosis-indexed. Cohort-specific follow-up timing is detailed in Supplementary Table 20. SCAN-B OS was diagnosis-indexed, and the original release excluded post-neoadjuvant specimens.

## Ethics statement

This study was a secondary analysis of public, de-identified cancer datasets and involved no new participant recruitment, intervention or specimen collection. No institutional patient data were included. Source-study ethics and consent information accompanies the original data releases.

## Data availability

TCGA molecular data are publicly available through UCSC Xena [30], and overall-survival endpoints through the TCGA Pan-Cancer Clinical Data Resource [29]. Open-access H&E slides and metadata are available through the GDC Data Portal [32]. CPTAC processed RNA expression data were obtained from LinkedOmics [38] and the CPTAC pan-cancer release [21]; clinical records were obtained from the corresponding releases, cBioPortal [39] and GDC, as specified in Methods. SCAN-B RNA expression and clinical records were obtained from the original GSE96058 release in the Gene Expression Omnibus (GEO) [22]; adopted versions, access URLs and OS definitions for the external cohorts are listed in Supplementary Table 20.

## Code availability

The source code for this study will be made publicly available upon publication.

## Acknowledgments

We thank the TCGA Research Network, the Clinical Proteomic Tumor Analysis Consortium (CPTAC), and the SCAN-B investigators and study participants for making their data available. Computational resources were provided by the Ohio Supercomputer Center. ChatGPT was used to improve manuscript clarity and formatting. The authors reviewed the content and take full responsibility for it.

## Author contributions

P.L., Z.M., M.H. and E.H. contributed to study conception and design. P.L. developed the methodology and software, curated the data, performed the analyses and visualization, and wrote the original draft. Z.M., M.K.K.N., M.H. and E.H. contributed to interpretation of the results and manuscript review and editing. M.H. and E.H. acquired funding. E.H. supervised the study.

---

**Extended Data Fig. 1.**
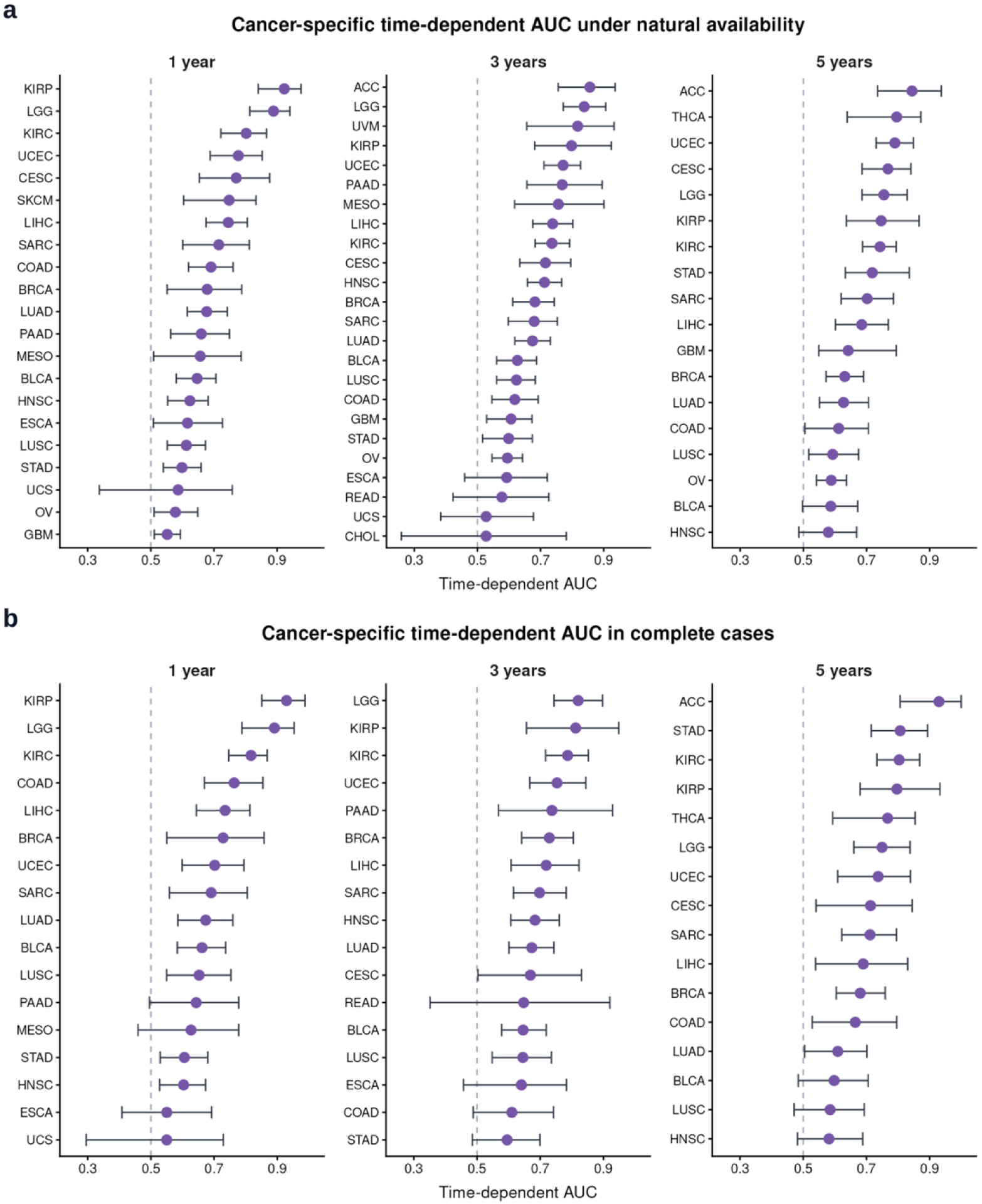
Cancer-specific time-dependent discrimination. **a,b**, Cumulative/dynamic AUC at 1, 3 and 5 years, estimated using inverse probability of censoring weights under natural availability (a) and in complete cases (b). Points are five-seed means of fold-aggregated AUCs; bars are percentile 95% intervals from 1,000 patient bootstrap samples. Cancer–horizon combinations are displayed only when minimum case, control and evaluable-fold requirements are met (Methods); statuses and counts are reported in Supplementary Table 7. Dashed lines denote AUC = 0.5.

**Extended Data Fig. 2.**
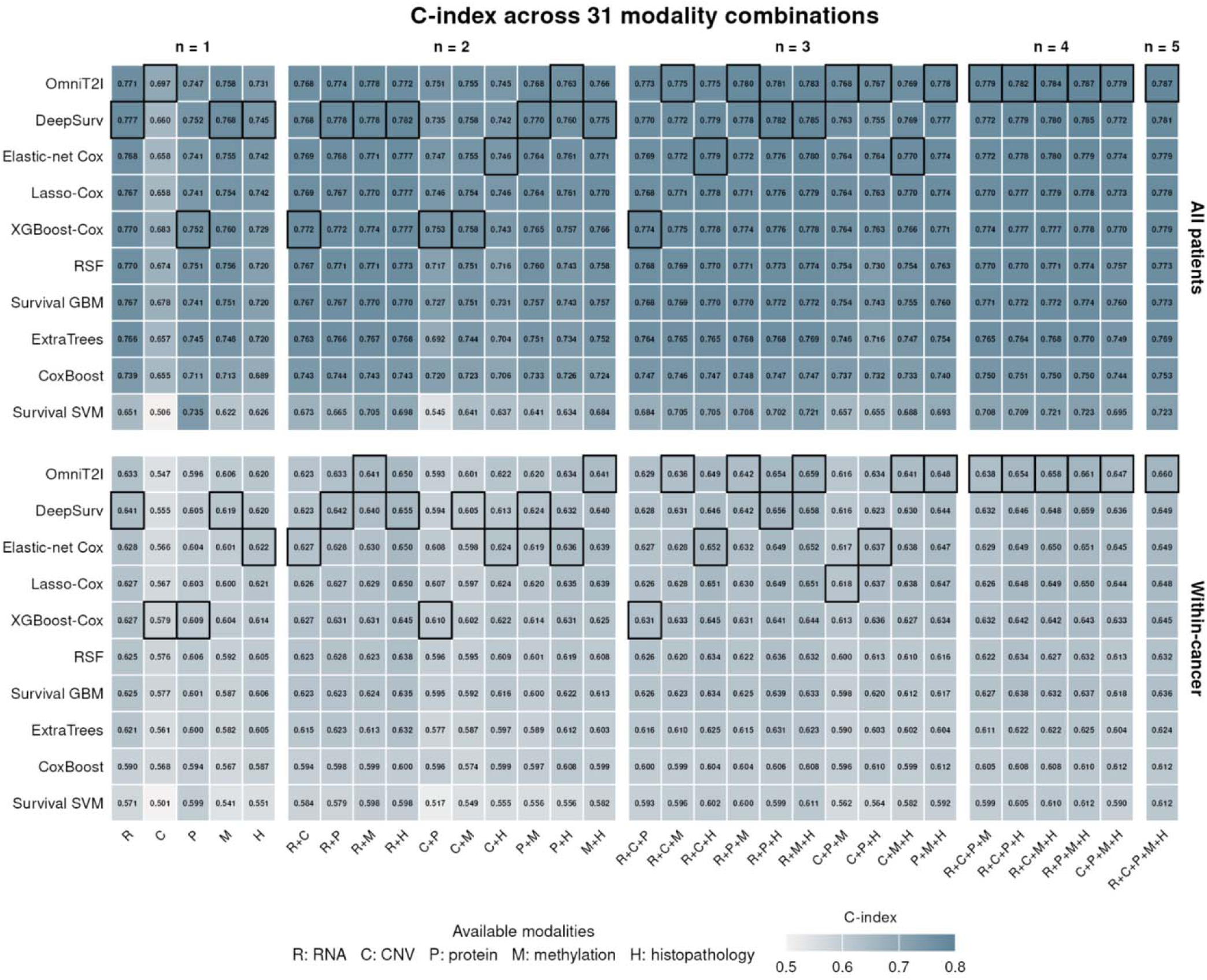
Absolute discrimination across all 31 measured-modality combinations. Pooled Harrell C-index (top) and within-cancer C-index (bottom). The rows include OmniT2I and nine specialist families. Each cell is the five-seed mean for one model and modality subset, evaluated using out-of-fold predictions for the same 5,605 complete-case patients (1,546 deaths). Black outlines identify the largest point estimate in each column. Values are printed to three decimals; color saturation spans 0.50–0.80. R, RNA; C, CNV; P, protein; M, methylation; H, histopathology.

**Extended Data Fig. 3.**
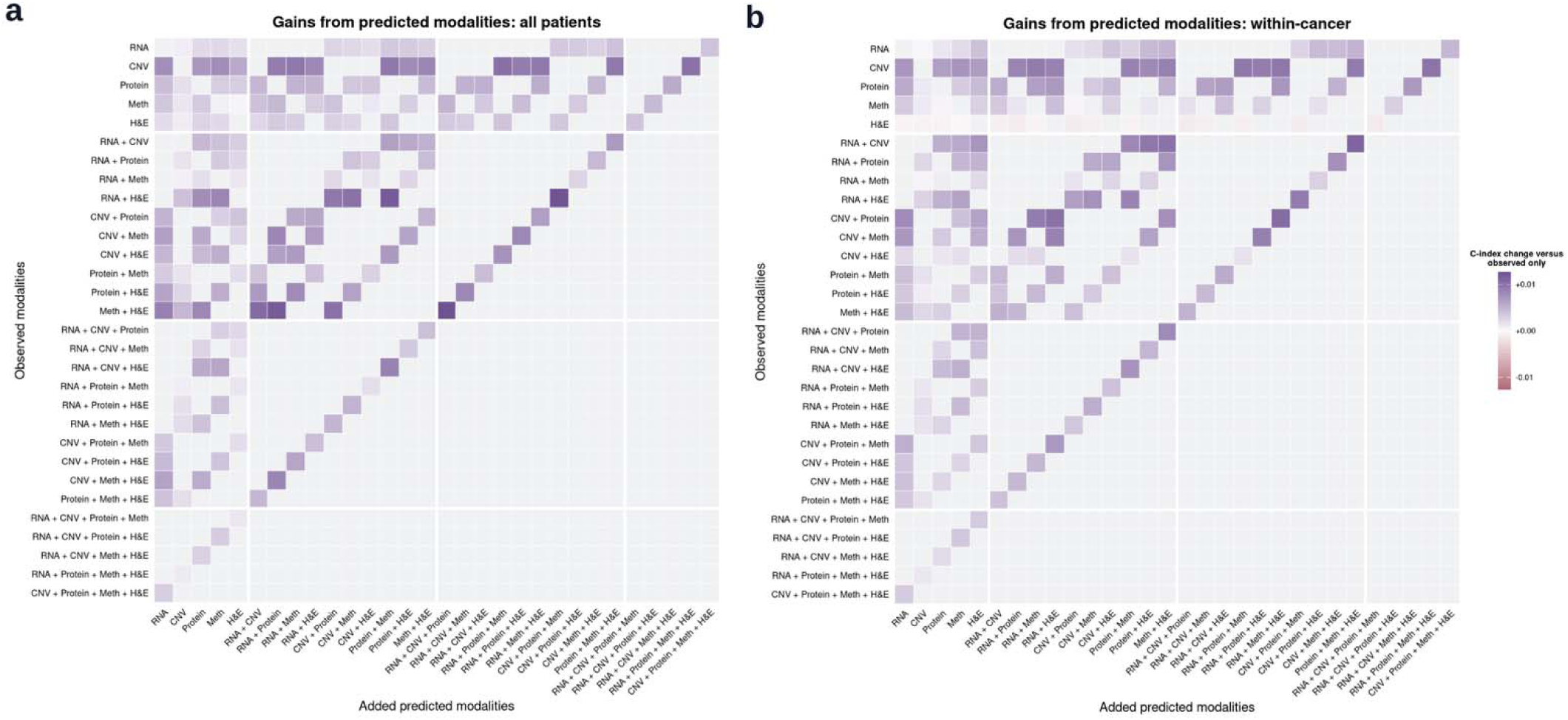
Gains from predicted modalities across all 180 eligible source–target routes. **a**, Pooled C-index. **b**, Within-cancer C-index. Each cell shows the five-seed mean paired difference between source plus predicted target modalities and the same measured sources alone, on 5,605 complete-case patients (1,546 deaths). Rows identify observed source subsets; columns identify added predicted target subsets. Source and target sets are nonempty and disjoint. Gray cells are ineligible routes, not zero-valued effects. Purple denotes positive and rose negative differences, with shared symmetric color limits. There are 180 positive pooled and 164 positive within-cancer differences.

## References

1. Hoadley, K. A. et al. Cell-of-origin patterns dominate the molecular classification of 10,000 tumors from 33 types of cancer. Cell 173, 291–304.e6 (2018). 10.1016/j.cell.2018.03.022

2. Cheerla, A. & Gevaert, O. Deep learning with multimodal representation for pancancer prognosis prediction. Bioinformatics 35, i446–i454 (2019). 10.1093/bioinformatics/btz342

3. Chen, R. J. et al. Pan-cancer integrative histology-genomic analysis via multimodal deep learning. Cancer Cell 40, 865–878.e6 (2022). 10.1016/j.ccell.2022.07.004

4. Fan, Z., Jiang, Z., Liang, H. & Han, C. Pancancer survival prediction using a deep learning architecture with multimodal representation and integration. Bioinform. Adv. 3, vbad006 (2023). 10.1093/bioadv/vbad006

5. Chen, R. J. et al. Multimodal Co-Attention Transformer for Survival Prediction in Gigapixel Whole Slide Images. In Proceedings of the IEEE/CVF International Conference on Computer Vision 3995–4005 (2021). 10.1109/ICCV48922.2021.00398

6. Jaume, G. et al. Modeling Dense Multimodal Interactions Between Biological Pathways and Histology for Survival Prediction. In Proceedings of the IEEE/CVF Conference on Computer Vision and Pattern Recognition 11579–11590 (2024). 10.1109/CVPR52733.2024.01100

7. Vale-Silva, L. A. & Rohr, K. Long-term cancer survival prediction using multimodal deep learning. Sci. Rep. 11, 13505 (2021). 10.1038/s41598-021-92799-4

8. Song, Y. et al. A cancer-type-aware framework for robust multimodal survival prediction under missing modalities. Brief. Bioinform. 27, bbag124 (2026). 10.1093/bib/bbag124

9. Zanitti, M. et al. A contrastive variational autoencoder for NSCLC survival prediction with missing modalities. In 2025 IEEE International Conference on Big Data (BigData) 2074–2083 (IEEE, 2025). 10.1109/BigData66926.2025.11401431

10. Ma, S. et al. Moving towards genome-wide data integration for patient stratification with Integrate Any Omics. *Nat*. Mach. Intell. 7, 29–42 (2025). 10.1038/s42256-024-00942-3

11. Cohen Kalafut, N., Huang, X. & Wang, D. Joint variational autoencoders for multimodal imputation and embedding. *Nat*. Mach. Intell. 5, 631–642 (2023). 10.1038/s42256-023-00663-z

12. Wissel, D. et al. SurvBoard: standardized benchmarking for multi-omics cancer survival models. Brief. Bioinform. 26, bbaf521 (2025). 10.1093/bib/bbaf521

13. Sharma, A., Vans, E., Shigemizu, D., Boroevich, K. A. & Tsunoda, T. DeepInsight: a methodology to transform a non-image data to an image for convolution neural network architecture. Sci. Rep. 9, 11399 (2019). 10.1038/s41598-019-47765-6

14. Ma, S. & Zhang, Z. OmicsMapNet: transforming omics data to take advantage of deep convolutional neural network for discovery. Preprint at 10.48550/arXiv.1804.05283 (2018).

15. Zhu, Y. et al. Converting tabular data into images for deep learning with convolutional neural networks. Sci. Rep. 11, 11325 (2021). 10.1038/s41598-021-90923-y

16. Tang, X. et al. OmicsFootPrint: a framework to integrate and interpret multi-omics data using circular images and deep neural networks. Nucleic Acids Res. 52, e99 (2024). 10.1093/nar/gkae915

17. Kalakoti, Y., Yadav, S. & Sundar, D. SurvCNN: a discrete time-to-event cancer survival estimation framework using image representations of omics data. Cancers 13, 3106 (2021). 10.3390/cancers13133106

18. Jiang, Y. et al. An interpretable omics-to-image transformer framework for cancer prognosis prediction. Comput. Struct. Biotechnol. J. 35, 0069 (2026). 10.34133/csbj.0069

19. McGough, S. F. et al. Prognostic pan-cancer and single-cancer models: A large-scale analysis using a real-world clinico-genomic database. PLoS ONE 21, e0341355 (2026). 10.1371/journal.pone.0341355

20. Xu, H. et al. A whole-slide foundation model for digital pathology from real-world data. Nature 630, 181–188 (2024). 10.1038/s41586-024-07441-w

21. Li, Y. et al. Proteogenomic data and resources for pan-cancer analysis. Cancer Cell 41, 1397–1406 (2023). 10.1016/j.ccell.2023.06.009

22. Brueffer, C. et al. Clinical Value of RNA Sequencing–Based Classifiers for Prediction of the Five Conventional Breast Cancer Biomarkers: A Report From the Population-Based Multicenter Sweden Cancerome Analysis Network—Breast Initiative. JCO Precis. Oncol. 2, 1–18 (2018). 10.1200/PO.17.00135

23. Huang, C. et al. Proteogenomic insights into the biology and treatment of HPV-negative head and neck squamous cell carcinoma. Cancer Cell 39, 361–379.e16 (2021). 10.1016/j.ccell.2020.12.007

24. Yuan, W. et al. Pancancer outcome prediction via a unified weakly supervised deep learning model. Signal Transduct. Target. Ther. 10, 285 (2025). 10.1038/s41392-025-02374-w

25. Zhou, F. & Chen, H. Cross-Modal Translation and Alignment for Survival Analysis. In Proceedings of the IEEE/CVF International Conference on Computer Vision 21428–21437 (2023). 10.1109/ICCV51070.2023.01964

26. Robinet, L., Berjaoui, A. & Cohen-Jonathan Moyal, E. Masked omics modeling for multimodal representation learning across histopathology and molecular profiles. Preprint at 10.48550/arXiv.2508.00969 (2025).

27. Wu, K. E., Yost, K. E., Chang, H. Y. & Zou, J. BABEL enables cross-modality translation between multiomic profiles at single-cell resolution. Proc. Natl Acad. Sci. USA 118, e2023070118 (2021). 10.1073/pnas.2023070118

28. Nikolaou, N. et al. A machine learning approach for multimodal data fusion for survival prediction in cancer patients. *npj Precis*. Oncol. 9, 128 (2025). 10.1038/s41698-025-00917-6

29. Liu, J. et al. An integrated TCGA pan-cancer clinical data resource to drive high-quality survival outcome analytics. Cell 173, 400–416.e11 (2018). 10.1016/j.cell.2018.02.052

30. Goldman, M. J. et al. Visualizing and interpreting cancer genomics data via the Xena platform. Nat. Biotechnol. 38, 675–678 (2020). 10.1038/s41587-020-0546-8

31. Mermel, C. H. et al. GISTIC2.0 facilitates sensitive and confident localization of the targets of focal somatic copy-number alteration in human cancers. Genome Biol. 12, R41 (2011). 10.1186/gb-2011-12-4-r41

32. Heath, A. P. et al. The NCI Genomic Data Commons. Nat. Genet. 53, 257–262 (2021). 10.1038/s41588-021-00791-5

33. Harrell, F. E. Jr., Califf, R. M., Pryor, D. B., Lee, K. L. & Rosati, R. A. Evaluating the yield of medical tests. JAMA 247, 2543–2546 (1982). 10.1001/jama.1982.03320430047030

34. Uno, H., Cai, T., Tian, L. & Wei, L. J. Evaluating prediction rules for t-year survivors with censored regression models. J. Am. Stat. Assoc. 102, 527–537 (2007). 10.1198/016214507000000149

35. Pölsterl, S. scikit-survival: A Library for Time-to-Event Analysis Built on Top of scikit-learn. J. Mach. Learn. Res. 21, 1–6 (2020). https://www.jmlr.org/papers/v21/20-729.html

36. Katzman, J. L. et al. DeepSurv: personalized treatment recommender system using a Cox proportional hazards deep neural network. BMC Med. Res. Methodol. 18, 24 (2018). 10.1186/s12874-018-0482-1

37. Chen, T. & Guestrin, C. XGBoost: A Scalable Tree Boosting System. In Proceedings of the 22nd ACM SIGKDD International Conference on Knowledge Discovery and Data Mining 785–794 (ACM, 2016). 10.1145/2939672.2939785

38. Vasaikar, S. V., Straub, P., Wang, J. & Zhang, B. LinkedOmics: analyzing multi-omics data within and across 32 cancer types. Nucleic Acids Res. 46, D956–D963 (2018). 10.1093/nar/gkx1090

39. Cerami, E. et al. The cBio cancer genomics portal: an open platform for exploring multidimensional cancer genomics data. Cancer Discov. 2, 401–404 (2012). 10.1158/2159-8290.CD-12-0095

